# Myofibrillar protein accumulation but reduced protein synthesis in PDCD4-depleted myotubes

**DOI:** 10.1101/2025.10.24.684280

**Authors:** Stephen Mora, Lilia Alihemmat, Logan D. Davari, Termeh Ataie, Parsa Nafari Mehranjani, Luke Flewwelling, Arthur J. Cheng, Olasunkanmi A.J. Adegoke

## Abstract

Skeletal muscle is critical to whole-body functionality and homeostasis. The mammalian/mechanistic target of rapamycin complex 1 (mTORC1) is a nutrient/growth-factor sensitive positive regulator of skeletal muscle mass. Amongst other substrates, mTORC1 phosphorylates the ribosomal protein S6 kinase (S6K1). Activated S6K1 acts through multiple effectors, including programmed cell death 4 (PDCD4), to activate mRNA translation and protein synthesis. Much of what is known about PDCD4 is in non-muscle cells. We previously demonstrated that the effect of PDCD4 differs between myoblasts and myotubes. Here, we showed that PDCD4 depletion in L6 and C2C12 myotubes enhanced myotube diameter (+36%) and accumulation of myofibrillar proteins (+163 – 237%). These effects occurred along with increased phosphorylation of AKT^ser473^ (+85%) and of the mTORC1 substrate S6K1^thr389^ (+152%), but protein synthesis was suppressed. There was increased phosphorylation of FoxO3a^ser253^ (+250%) and a corresponding reduction in the expression of the muscle protein ubiquitin ligase MuRF1 (–44%), but there was no significant effect on measures of proteolysis or autophagy. In starved myotubes treated with the proteasome inhibitor MG132, accumulation of ubiquitinated proteins was attenuated in PDCD4-depleted cells. PDCD4 depletion did not augment measures of myotube contraction but was associated with reduced ATP and intracellular amino acid levels. Finally, AKT inhibition partially attenuated the effect of PDCD4 depletion on myofibrillar protein abundance. In summary, myofibrillar protein accumulation in PDCD 4-depleted myotubes did not lead to improved myotube function, likely due to reduced energy level. Our data point to a pivotal role for PDCD4 in regulating myotube size and metabolism.

## Introduction

The preservation of skeletal muscle mass and metabolism is essential for locomotion (1), vital organ function (2) and whole-body substrate homeostasis (3). As such, loss of skeletal muscle mass/function can lead to or worsen chronic metabolic diseases, such as obesity, diabetes, and cancer (4, 5). Interventions that minimize the loss of muscle mass/function could have therapeutic potential in the management of these diseases.

Muscle mass is determined in part by protein mass, which is the net sum of the rates of muscle protein synthesis and proteolysis. Critical in the regulation of skeletal muscle mass is the growth factor/nutrient-sensitive kinase complex, the mammalian/mechanistic target of rapamycin complex 1 (mTORC1) (6, 7). Upstream, mTORC1 is activated by pro-anabolic signals, such as growth factors (8), amino acids (9), glucose (10), oxygen (11) and energy levels (12). When activated, mTORC1 phosphorylates, among other substrates, two regulators of mRNA translation and ribosomal biogenesis: ribosomal S6 protein kinase 1 (S6K1) and the eukaryotic mRNA translation initiation factor (eIF) 4E-binding protein 1 (4E-BP1) (6, 13). S6K1 promotes skeletal muscle protein synthesis through the phosphorylation of ribosomal protein S6 (S6) (14) and eIF4B (15), whereas 4E-BP1 is an inhibitor of cap-dependent mRNA translation that is inhibited by mTORC1-catalyzed phosphorylation. There is evidence to suggest that substrates other than S6, including programmed cell death protein 4 (PDCD4), are critical to the effects of mTORC1/S6K1 on muscle anabolism (16). In addition to protein synthesis, mTORC1 can also regulate proteolysis via its effect on autophagy (17) and possibly the ubiquitin dependent proteolytic system (18, 19).

PDCD4 is an mRNA translation inhibitor (20), tumour suppressor (20) and cell cycle inhibitor (21). Loss of PDCD4 is associated with the progression of colon (22), breast (23), and lung (24) carcinomas. In the unphosphorylated state, PDCD4 inhibits mRNA translation by binding to eIF4A and eIF4G, two factors critical for the formation of a complex that is required for mRNA translation initiation, the eIF4F complex (25). Activated S6K1 phosphorylates PDCD4^Ser67^ leading to the ubiquitination of PDCD4 by beta-transducing repeat-containing protein (βTrCP) ubiquitin ligase, followed by its subsequent degradation by the proteasome. This permits initiation of mRNA translation (16).

Not much is known about the role of PDCD4 in skeletal muscle. Previously, we showed that PDCD4 levels in skeletal muscle are sensitive to nutritional manipulations (26) and that myoblasts depleted of PDCD4 are impaired in their ability to form myotubes (27). We have also shown that the effect of PDCD4 on protein synthesis is dependent on the nature of the cells: PDCD4 depletion increases protein synthesis in myoblasts, but not in myotubes (28). However, whether PDCD4 regulates myofibrillar protein abundance or other aspects of myotube protein metabolism has not been addressed. Given what is known about PDCD4 as an inhibitor of mRNA translation initiation, we hypothesized that PDCD4 depletion in myotubes would augment myotube protein abundance mediated at least in part by reduced proteolysis. Here, we showed that PDCD4 depletion improved myotube size, increased myofibrillar protein abundance and AKT phosphorylation (activation). Surprisingly, these changes occurred along with reduced protein synthesis, minimal effects on proteolysis, and reduced myotube free amino acid and ATP levels. We also showed that inhibiting AKT activation partially attenuated the effect of PDCD4 depletion on myofibrillar protein abundance.

## Materials and Methods

### Antibodies

Antibodies to PDCD4 (#9535), p62 (#5114), beclin-1 (#3738), microtubule-associated proteins 1A/1B light chain 3B (LC3B) (#3868), phosphorylated (p) FoxO3a^ser253^ (#9466), p-AKT^Ser473^ (#4060), p-S6^Ser235/236^ (#4858), and its kinase S6K1^thr389^ (#9234) were purchased from Cell Signalling Technology (Danvers, MA). Antibodies to tropomyosin (CH1), troponin (JLT12) and myosin heavy chain-1 (MHC-1)(MF-20) were purchased from Developmental Hybridoma (Iowa City, Iowa, USA). Antibody against the ubiquitin protein ligase muscle RING-finger protein-1 (MuRF1, #55456-1-AP) was obtained from Protein Tech (San Diego, CA, USA). Anti-ubiquitin antibody (#sc-8017) was purchased from Santa Cruz Technology (Dallas, TX, USA) while antibodies against γ-tubulin (#T6557), and puromycin (#MABE343) were purchased from Sigma Aldrich (St. Louis, MO, USA). We purchased ^14^C-valine (#ARC-0678) from American Radiolabeled Chemicals Inc. (St Lous, MO, USA).

### Cell Culture

L6 myoblasts were cultured in 6 or 12-well plates (depending on the question being asked) in growth medium (GM: AMEM supplemented with 10% FBS and 1% antibiotic-antimycotic reagents) at 37°C and 5% CO_2_. To induce myotube differentiation, confluent cells were shifted into differentiation medium (DM: AMEM supplemented with 1% antibiotic-antimycotic reagents and 2% horse serum). Medium was replaced every other day until myotubes were formed. For C2C12 experiments, myoblasts were grown at 37°C and 5% CO_2_ in DMEM supplemented with 10% FBS and 1% antibiotic-antimycotic reagents. Once confluent, cells were shifted into DMEM supplemented with 1% antibiotic-antimycotic reagents and 2% horse serum.

### Gene Knockdown

RNA interference (RNAi) was done as previously described (28). Myotubes were incubated in either control (scrambled) or siRNA (designed against genes of interest) oligonucleotide transfection media, using lipofectamine RNAiMAX (Thermo Fisher, #2179825) following manufacturer’s instructions. Control transfection medium consisted of scramble oligonucleotides (Sigma Aldrich, #SIC001), Opti-MEM (Gibco, #31985-070) and 1mL of AMEM supplemented with 10% FBS (antibiotic-antimycotic free). We used the following PDCD4 siRNA oligonucleotides, all from Millipore Sigma, Oakville ON Canada: PDCD4 #1 sense (GUCUUCUACUAU UACCAUA [dT] [dT]), PDCD4 #1 antisense (5′UAUGG UAAUAGUAGAAGAC [dT] [dT]), PDCD4 #2 sense (CUACUAUUACCAUAGACCA [dT] [dT]), and PDCD4 #2 antisense (UGGUCUAUGGUAAUAGUAG [dT] [dT]. Twenty-four hours following transfection, 1mL of AMEM supplemented with 10% FBS and 1% antibiotic-antimycotic was added to each well. Forty-eight hours following transfection, cells were washed 2X in phosphate-buffered-saline (PBS) and harvested with 100µL of lysis buffer (final concentration: 1mM ethylenediaminetetraacetic acid (EDTA), 2% sodium dodecyl sulfate (SDS), 25 mM Tris-HCl pH 7.5, 10 μL/mL protease inhibitor cocktail (Sigma Aldrich, #P8340-5ML),10 μL/mL phosphatase inhibitor cocktail (Sigma Aldrich, #P5726-5ML), 1 mM dithiothreitol (DTT). Sample lysates were stored at -20°C until analysis. Western blotting was used to confirm successful knockdown of PDCD4 in myotubes. We routinely observed > 90% knockdown efficiency (Fig. 1A). This transfection protocol was carried out similarly for C2C12 myotubes using oligonucleotides designed against mouse PDCD4.

**Fig 1.**
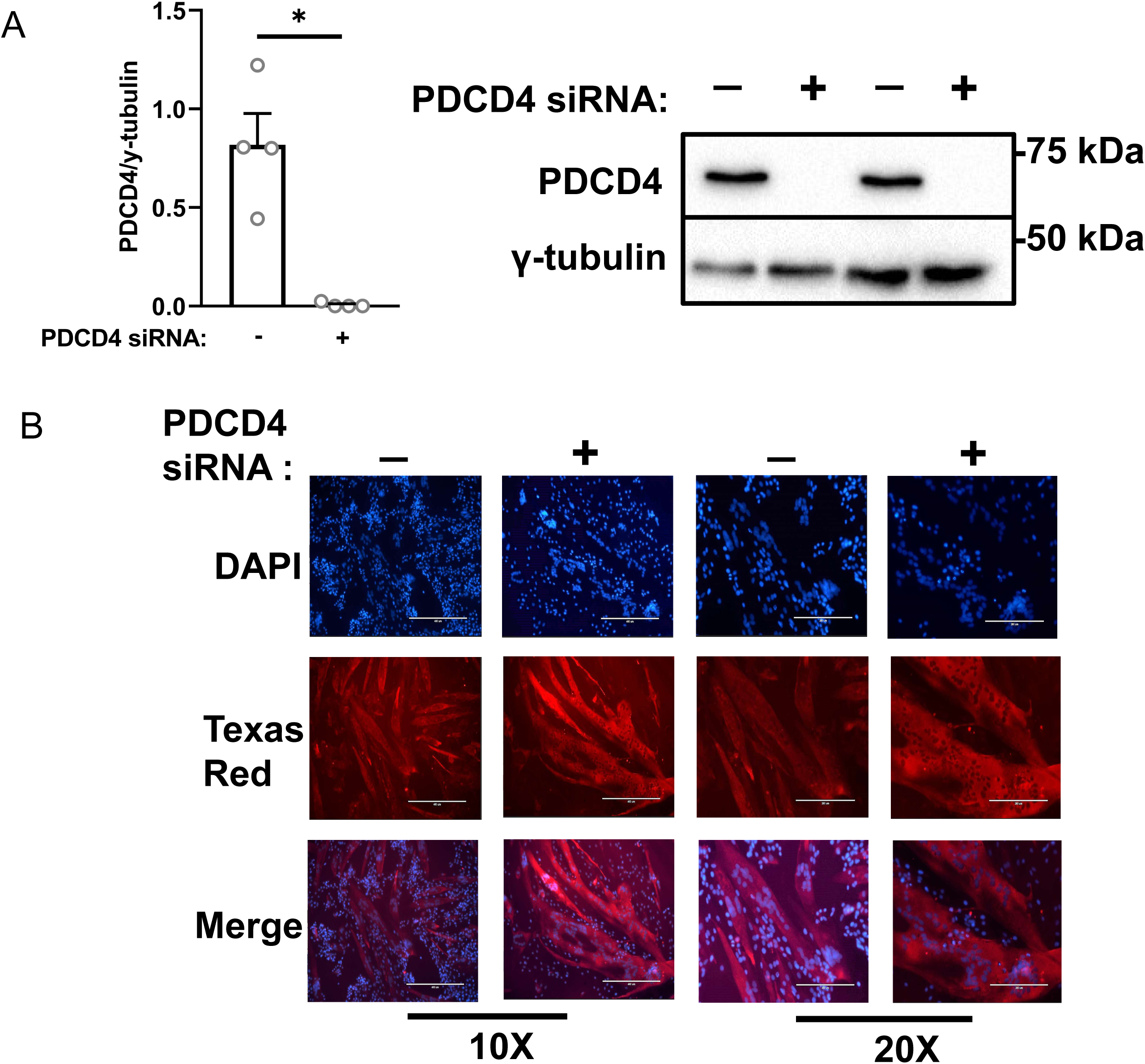
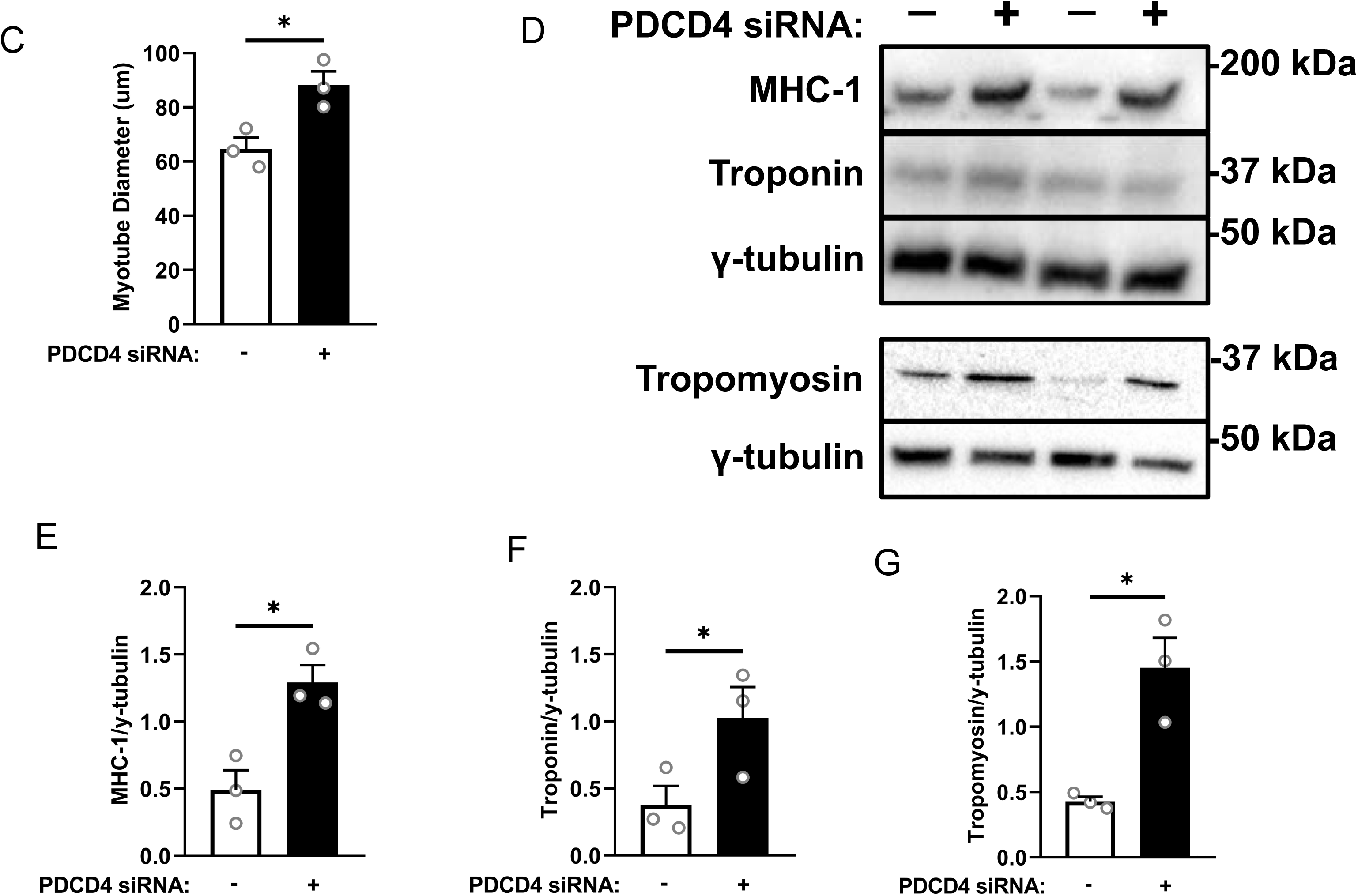
PDCD4 depletion in L6 myotubes enhances myofibrillar protein abundance. A: Myotubes were treated with either scrambled (SCR, white bars, ‘-‘) or PDCD4 (black bars, ‘+’) siRNA oligonucleotides. Successful depletion of PDCD4 is shown. B: Immunofluorescence detection of MHC-1 and nuclei at 10x (Bar, 400µM) and 20x (Bar, 200µM) objective. C: Quantified myotube diameter (μm). Immunoblotting (D) and quantified data for MHC-1 (E), troponin (F) and tropomyosin (G). Data are mean ± SEM, n=3-4 independent experiments with 3 technical replicates per experiment, * p < 0.05.

### Immunofluorescence Microscopy

Scramble and PDCD4-depleted myotubes cultured on cover slips (Fisher Scientific, #092815-9) within 6-well plates were fixed (4% paraformaldehyde in PBS, 10 min, room temp), permeabilized (0.03% Triton X-100 in PBS, 5 min, room temp) and blocked (10% horse serum in PBS) for 1h at 37°C. Myotubes were then incubated in primary antibody (2.5μg/mL of anti MHC-1 antibody) in 1% bovine serum albumin (BSA)) overnight at 4°C. The following day, myotubes were washed 3 X 5 minutes in PBS at room temperature and incubated in secondary antibody (Texas Red anti-mouse IgG secondary antibody (1:100 with 1% BSA)) for 2h at room temperature. Next, 4′,6-diamidino-2-phenylindole (DAPI) staining (for nuclei) and mounting the cover slip on a microscope slide were completed. Slides were then imaged using the EVOS FL Auto microscope (Life Technologies) along with the EVOS FL Auto program for maintaining acquisition settings (light and brightness) in all experimental treatments as described previously (29, 30). Briefly, all images were transformed into an 8-bit gray scale image and the mean gray value of each sample was measured within a 0–255 range using ImageJ. Images were quantified using ImageJ.

### Protein Synthesis (Sunset Analysis)

Control and PDCD4-depleted myotubes were starved (serum-free, amino acid-free medium, US Biological, #R8999-03) for 24h. They were then treated for 30 minutes with 1µM of puromycin diluted in differentiation medium as described before (31). Myotubes were then harvested in lysis buffer and proteins in the lysates were immunoblotted against an anti-puromycin antibody.

### Proteolysis

L6 myoblasts were seeded and differentiated in 6-well plates as described above. On day 4 of differentiation, myotubes were transfected with scrambled or PDCD4 siRNA oligonucleotides as described above. Immediately after adding the transfection mix, myotubes in each well were labeled with 5 μL of [¹ C]-valine (0.5 μCi/well) for 24 h. After, 1 mL of room-temperature differentiation media containing antibiotics was added to the existing media in each well, and the cells were incubated for an additional 24 h to allow further incorporation of the radiolabeled valine into proteins. After the 48-h labeling period, an aliquot of incubation medium was taken from each well and designated as medium radioactivity at the end of labelling period (or pulse period). A batch of wells was harvested (designated as time 0 below) while the remaining myotubes were rinsed 2X in PBS before being incubated in DM that was supplemented with 2mM non-radioactive Valine (‘chase’ solution) for 10 or 24h. At 0, 10 and 24 h of chase, samples were harvested by washing myotubes 2X in PBS and then incubating in 1 mL of cold 10% trichloroacetic acid (TCA, 15 min, on ice). Lysates were then collected into microcentrifuge tubes and kept on ice for 1 h. Samples were centrifuged at 9600xg for 2 minutes at 4°C, and the supernatant was discarded. Pellet was washed three times with cold 10% TCA. Following centrifugation (9600xg for 2 min at 4°C)), the pellet was resuspended in 300 μL of a solution of 0.1 M NaOH with 0.1% sodium deoxycholate. Following, the samples were neutralized with 1M HCl. An aliquot of this was counted in a scintillation counter. We calculated proteolysis by two approaches: *release* of ^14^C valine from previously labelled cellular proteins into the incubation medium, and by the *disappearance* of ^14^C-valine radioactivity from protein pellets at different times during the chase. The former was calculated by counting medium radioactivity at different times and expressing the data as a % of radioactivity in proteins at the end of the labelling period. The latter was derived by measuring residual radioactivity in proteins at different times and expressing the data as a % of radioactivity in proteins at the end of the labelling period.

### Starvation and Proteasome Inhibition

Myotubes were incubated in starvation medium as mentioned above for 24 hours. In the last 3h, starvation continued with or without the addition of the proteasome inhibitor MG132 (5µM). Myotubes were then harvested and immunoblotted for ubiquitin. Ponceau S staining was used to check equal protein loading.

### Electrically Induced Sarcoplasmic Reticulum Calcium Release (Myotube contraction)

C2C12 myoblasts were seeded and differentiated in 35 mm glass-bottom dishes (MatTek Corporation, Part No. P35G-1.5-14-C, Ashland, MA, USA) using DMEM-based DM. On day 4 of differentiation, myotubes were transfected with either PDCD4 siRNA oligonucleotides or scrambled control siRNA oligonucleotides using Lipofectamine RNAiMAX in Opti-MEM. Two days later, dishes were removed from the incubator, and excess media was removed from the glass-bottomed petri dish, leaving 2mL of media remaining in the dish. Myotubes were then loaded with [3.5 μM] of indo-1 AM (Thermo Fisher Cat # I1223) and placed in a water-saturated incubator (37 °C) for 30 min. Once finished, the dishes were moved to a dark room and placed into a custom-made 3D printed chamber and mounted on an inverted microscope (Nikon Eclipse Ts2R-FL, Tokyo, Japan) and continuously super-fused via an analog pump (Reglo, Ismatec, Glattburg, Switzerland) with room temperature (∼25°C) experimental Tyrode solution (mM): 121 NaCl, 5.0 KCl, 1.8 CaCl2, 0.5 MgCl_2_, 0.4 NaH_2_PO4, 24 NaHCO_3_, 0.1 EDTA, 5.5 glucose, and ∼0.2% fetal calf serum, bubbled with 95% O_2_/5% CO_2_ giving the bath a pH of ∼7.4. After one hour of washing the dishes, background fluorescence reading was acquired by focusing on an empty section of the dish. Myotubes were electrically stimulated in a custom-made chamber (holding two parallel platinum electrodes positioned on each side of the light path, 1 cm apart) connected to an electrical stimulator (Model 701C, Aurora Scientific, Aurora, ON, Canada) controlled via Aurora Scientific 600A software (Aurora Scientific, Aurora, ON, Canada). Felix32 software (Horiba, London, ON, Canada) was used to acquire cytoplasmic free [Ca^2+^] ([Ca^2+^]_i_) readings. Images were acquired using an inverted microscope equipped with a 40x objective. Indo-1 AM dye was excited at 346 ± 5 nm and the two emission wavelengths were recorded by two photomultipliers at 405±5 nm and 495±5 nm (Horiba Ratiomaster, London, ON, Canada) (32) Myotubes were stimulated at 200mA for 300ms at 100 Hz. Tetanic [Ca^2+^]_i_ was measured as the mean Indo-1 AM fluorescence ratio during tetanic stimulation trains. The size/area of the myotube(s) in the field of view was determined by exporting a screenshot of the viewed area to ImageJ, where the area was calculated. [Ca^2+^]_i_ was calculated per area of the myotube.

### Myotube ATP Assay

We used the ATP Determination Kit (Thermo Fisher Canada, Cat. # A22066) according to the manufacturer’s instructions. This bioluminescence-based assay uses firefly luciferase, which emits light in the presence of its substrate D-luciferin and ATP. L6 myotubes were transfected with either PDCD4 siRNA oligonucleotides or scrambled siRNA oligonucleotides as described before, and incubated for 48 hours prior to ATP analysis. Two days after transfection, cells were lysed and supernatants collected. A standard reaction solution was prepared by combining D-luciferin, recombinant firefly luciferase, reaction buffer, DTT, and deionized water, as per manufacturer’s instructions. A series of ATP standards were prepared to develop a standard curve. Samples and standards were added to the reaction solution in opaque white 96-well plates, with a final volume of 100 µL per well. Luminescence was measured immediately using a luminometer. Background signal was subtracted from all wells, and ATP concentrations were calculated by interpolating from the standard curve. Final values were normalized to total protein content (µmol ATP/g protein), as determined by the Pierce BCA Protein Assay Kit (Thermo Fisher, Cat #23225).

### Amino Acid Concentrations

This was done as described previously (33). L6 myotubes were transfected with control or PDCD4 siRNA oligonucleotides as described above. Two days later, myotubes were washed and harvested in 10% TCA. Cell lysates were then centrifuged at 2.3g for 15 minutes and the supernatant (containing free amino acids) was collected. The amino acids were then diluted in a ratio of 1 (sample): 2 (potassium borate buffer): 1 (0.1N hydrochloric acid): 8 (HPLC grade water). Diluted samples were pre-column derivatized with a ratio of 1 (sample): 1 (o-phthalaldehyde, 29.28mM). Samples were then injected into a YMC-Triart C18 column fitted onto an ultra HPLC system (Nexera X2, Shimadzu, Kyoto, Japan) connected to a fluorescence detector (Shimadzu, Kyoto, Japan; excitation: 340 nm; emission: 455 nm). A gradient solution derived from 20mM potassium phosphate buffer (pH 6.5) (Mobile phase A) and a solution made from HPLC grade water (15%), acetonitrile (45%) and methanol (40%) at a flow rate of 0.8mL/min was used to elute the amino acids. Amino acid standard curves were used to calculate amino acid concentrations and all samples were normalized to total protein as previously described.

### AKT Inhibition Experiments

Myotubes were treated with either scrambled or PDCD4 siRNA as described above. Twenty-four h after siRNA treatment, myotubes were treated with 5μM of an AKT inhibitor (AKT Inhibitor VIII, Millipore-Sigma Cat #124018). Myotubes were then harvested 24h following treatment with the AKT inhibitor.

### Western Blotting

Following harvesting, protein concentration was determined using the pierce BCA protein assay method (Thermo Fisher, Cat #23225). Approximately 25µg of protein (per sample) were separated on 10 or 15% SDS-PAGE gels. Proteins were then transferred onto polyvinylidene difluoride (PVDF) membranes (0.2µM, BIO-RAD). Primary and secondary antibody incubations, imaging and quantification have been described (30, 34).

### Statistical Analysis

All immunoblot analyses were quantified and adjusted to their corresponding γ-tubulin values, aside from ubiquitin, which was corrected to their respective ponceaus stain. Unpaired t-tests with a Welch correction were used to measure treatment differences between scrambled and PDCD4-depeleted myotubes. For starvation and proteasome inhibition experiments, a one-way ANOVA with a Tukey’s post-hoc test was used. Significance was determined when p-value < 0.05. Results were expressed as mean ± standard error of the mean (SEM) of at least 3 independent experiments (biological replicates, with 3 technical replicates per experiment).

## Results

### PDCD4 depletion in myotubes enhances myofibrillar protein abundance

To investigate the effect of PDCD4 in myotubes, we used siRNA to deplete this protein (Fig. 1A). PDCD4-depleted myotubes had larger diameters (Fig. 1B,C) and greater abundance of myofibrillar proteins MHC-1, troponin, and tropomyosin (Fig. 1D-G). Similar results were obtained in C2C12 myotubes depleted of PDCD4 (Fig. 2A – E).

**Fig 2.**
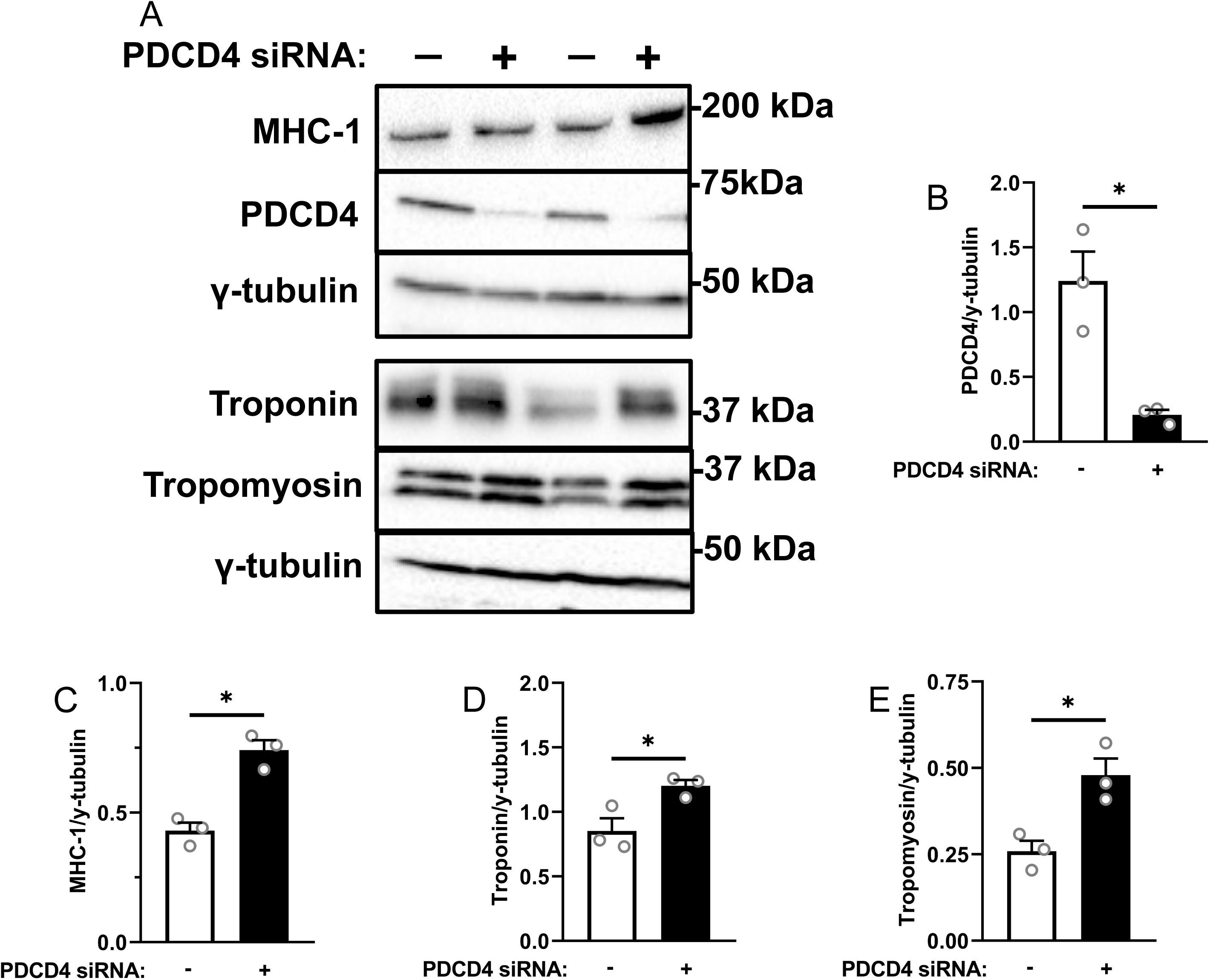
PDCD4 depletion in C2C12 myotubes enhances myofibrillar protein abundance. C2C12 myotubes were treated as described in figure 1 for L6 myotubes. Immunoblotting **(A)** and quantified data for PDCD4 **(B)**, MHC-1 **(C)**, troponin **(D)** and tropomyosin **(E).** Data are mean ± SEM, n=3 independent experiments, with 3 technical replicates per experiment, * p < 0.05.

### Anabolic signaling, but not protein synthesis, is enhanced in PDCD4-depleted myotubes

We investigated changes that might explain the increased abundance of myofibrillar proteins upon PDCD4 depletion, starting with mTORC1 signaling, a regulator of protein synthesis (35, 36). Phosphorylation of the upstream mTORC1 activator AKT^ser473^, was increased in PDCD4-depleted myotubes (Fig. 3A, B). Phosphorylation of a downstream target of mTORC1, S6K1^thr389^, was also increased (Fig. 3A, C). However, p-S6^ser235/236^, a S6K1 substrate, was not different between conditions (Fig. 3A, D). There was a trend for reduced 4E-BP1^thr37/46^ phosphorylation (Fig. 3A, E). The levels of unphosphorylated (total) AKT, S6K1 and S6 were not affected by treatments (Fig. 3F, G). Contrary to what one might expect from the increased abundance of myofibrillar proteins, but consistent with our previous findings (28), PDCD4 depletion resulted in a decrease in protein synthesis, as measured with sunset analysis (Fig. 3H).

**Fig 3.**
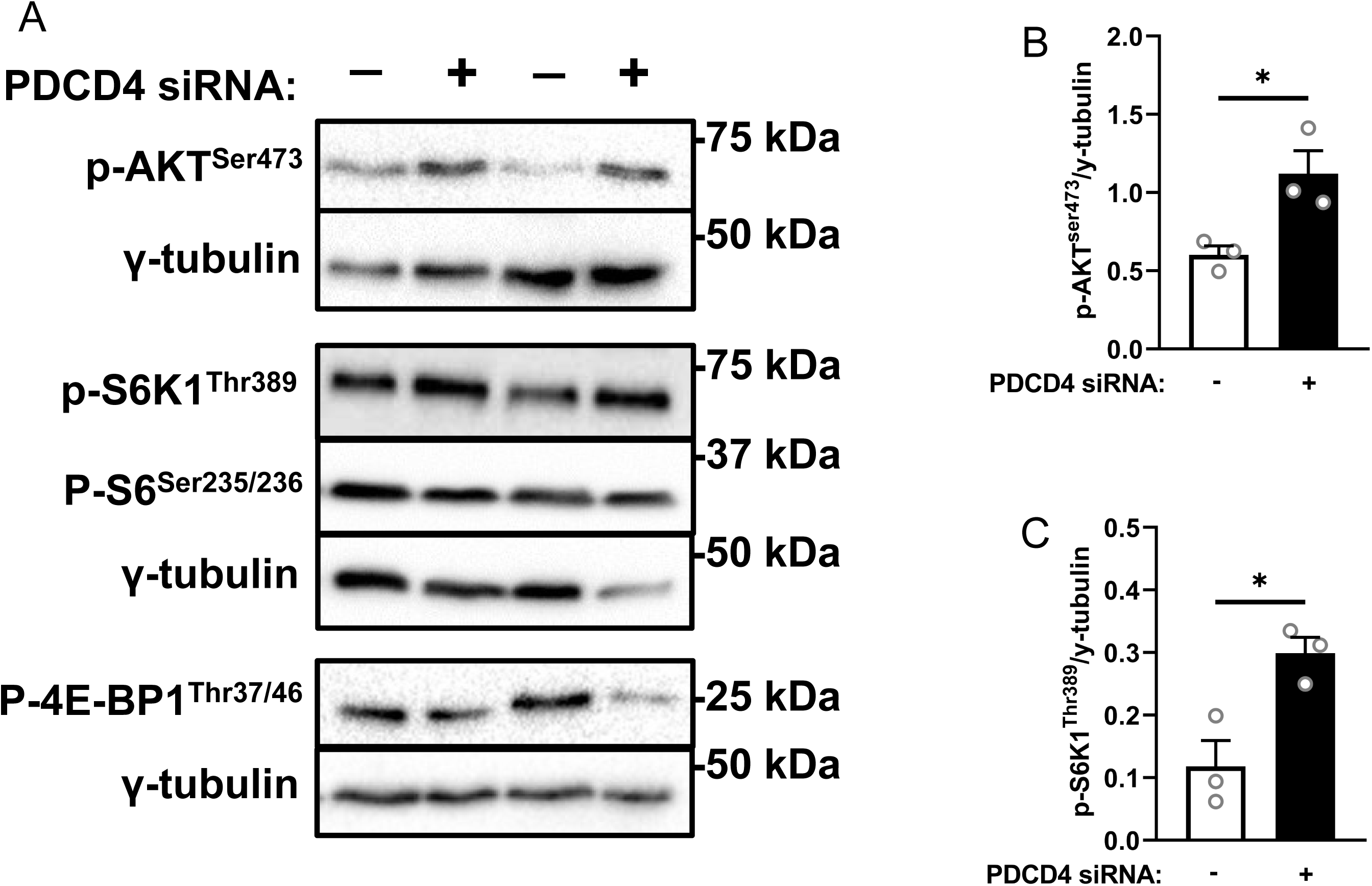
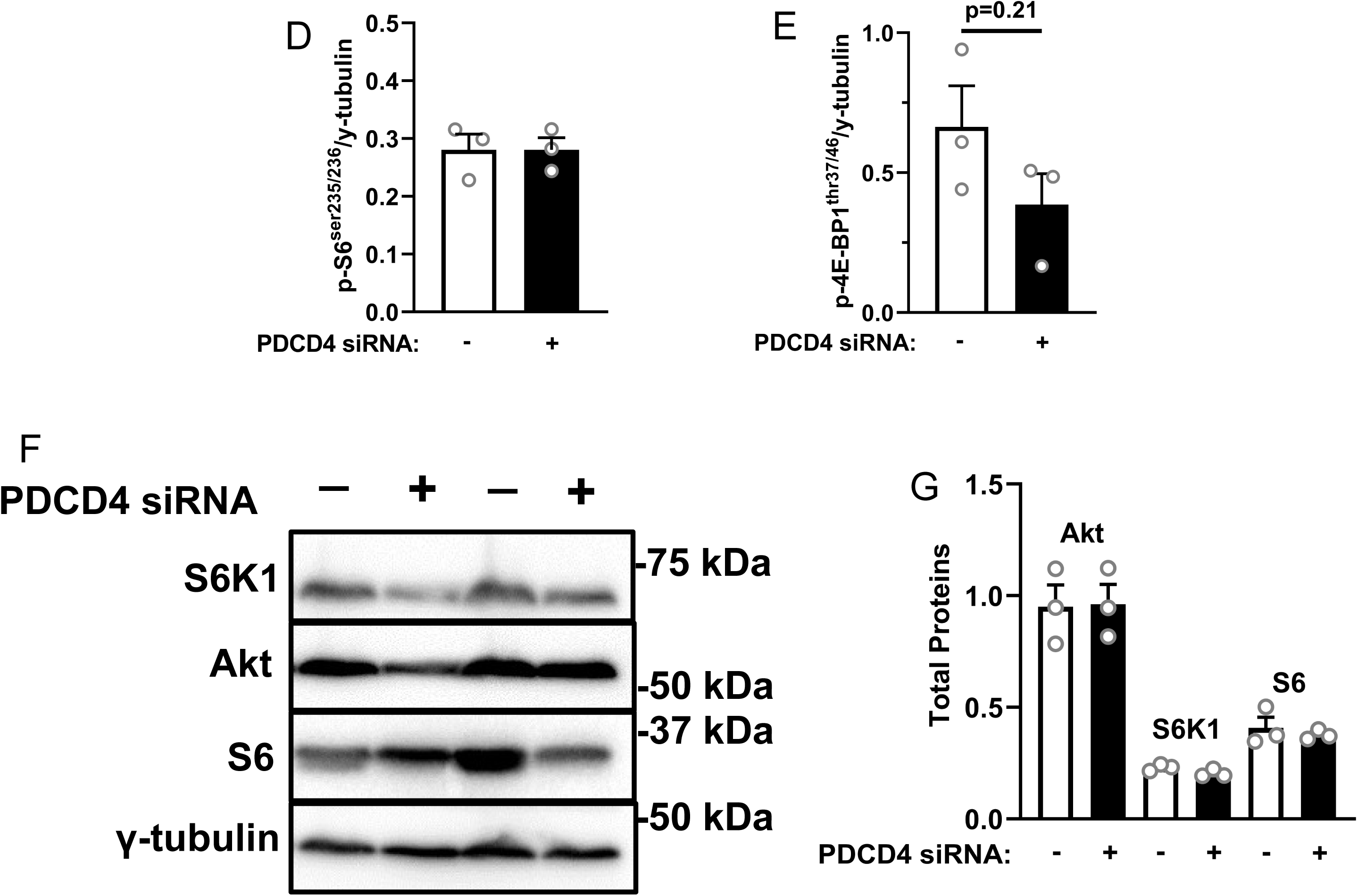
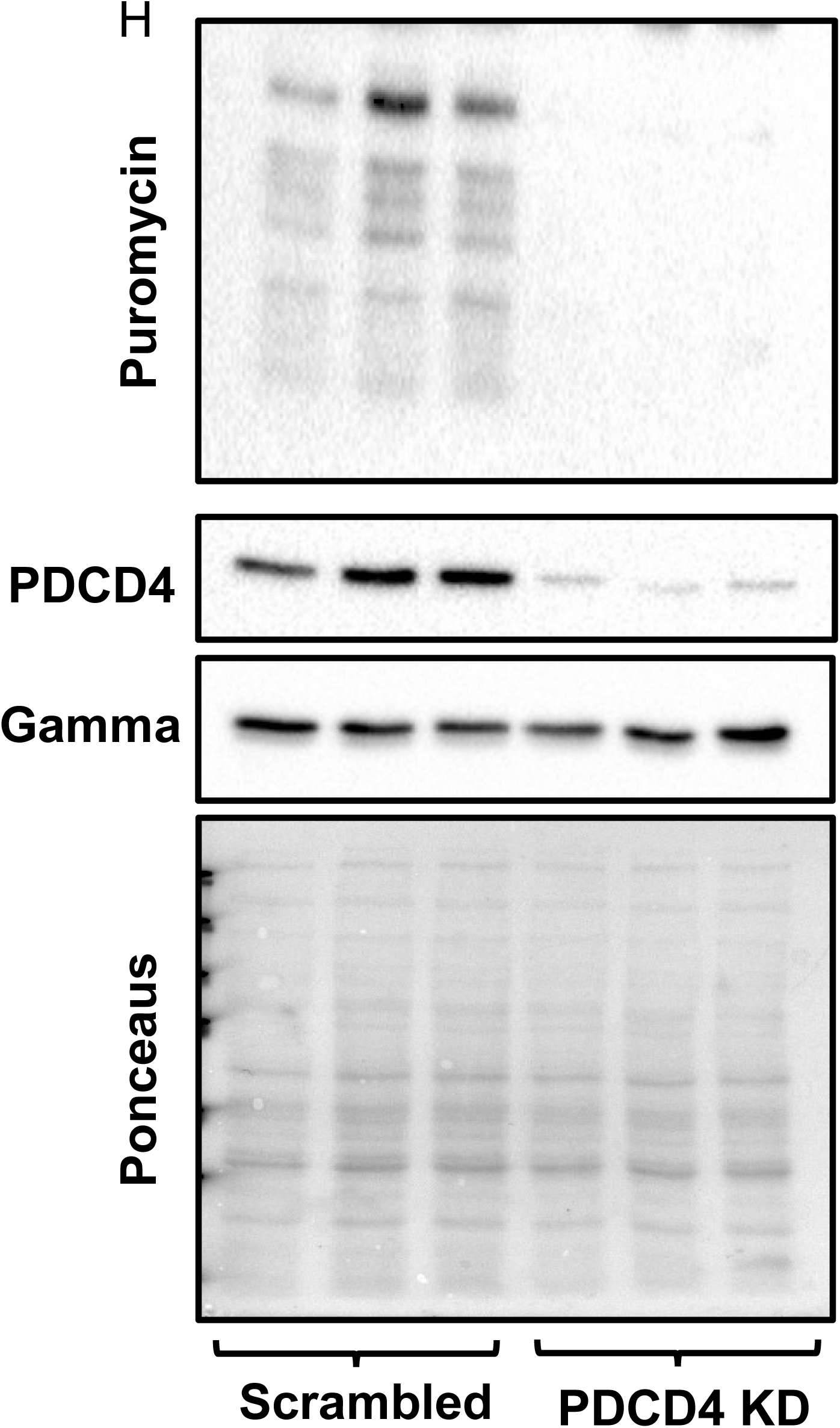
Anabolic signaling, but not protein synthesis, is enhanced in PDCD4-depleted myotubes. L6 myotubes were treated as described in figure 1. Immunoblotting **(A)** and quantified data for p-AKT^ser473^ **(B)**, S6K1^thr389^ **(C)**, S6^ser235/236^ **(D)** and 4E-BP1^thr37/46^ **(E)** are shown. Total protein for some of the above signals are shown **(F, G)**. Myotubes in the two treatment groups were incubated in 1µM of puromycin for 30 minutes and sunset analysis was performed **(H)**. Data are mean ± SEM, n=3 independent experiments with 3 technical replicates per experiment, * p < 0.05.

### PDCD4 depletion decreases MuRF1 protein abundance, but has no effect on proteolysis or measures of autophagy in myotubes

Increased anabolic signaling without a corresponding increase in protein synthesis led to us to hypothesize that myotubes depleted of PDCD4 would exhibit decreased protein catabolism. There was a trend toward increased FoxO3a^ser253^ phosphorylation (Fig. 4A, B). Consistent with this, PDCD4-depleted myotubes had reduced expression of the muscle ubiquitin protein ligase, MuRF1 (Fig. 4A, C). There was no significant treatment effect on the autophagy markers beclin-1, LC3BII/I, and p62 (Fig. 4A, D –F). To further probe treatment effect on proteolysis, we pre-labelled L6 myotube proteins with ^14^C-valine and then ‘chased’ for different lengths of time. The amount of radioactivity released into the incubation medium relative to total protein-associated radioactivity at the end of the incubation with ^14^C valine (‘pulse’) (positively proportional to the rate of proteolysis, Fig 4G) or the amount of ^14^C-radioactivity retained in myotube proteins relative to total protein-associated radioactivity at the end of the ‘pulse’ (inversely proportional to the rate of proteolysis) should each reflect proteolysis. PDCD4 knockdown did not significantly alter the rate of ^14^C-valine release from, or retention in myofibrillar proteins. To probe this further, we measured the abundance of ubiquitinated proteins, the substrates of the 26S proteasome. PDCD4 depletion had no effect on the abundance of ubiquitinated protein (Fig 4I, J). However, in cells treated with the proteasome inhibitor MG132, starvation-induced increase in the abundance of ubiquitinated proteins was attenuated in PDCD4-depleted cells (Fig. 4G, H). Taken together, these findings point to altered protein turnover in PDCD4-depleted cells with suppression of protein synthesis and minimal effect on proteolysis, although it is possible that proteolysis may be affected in altered physiological states (for example, starvation).

**Fig 4.**
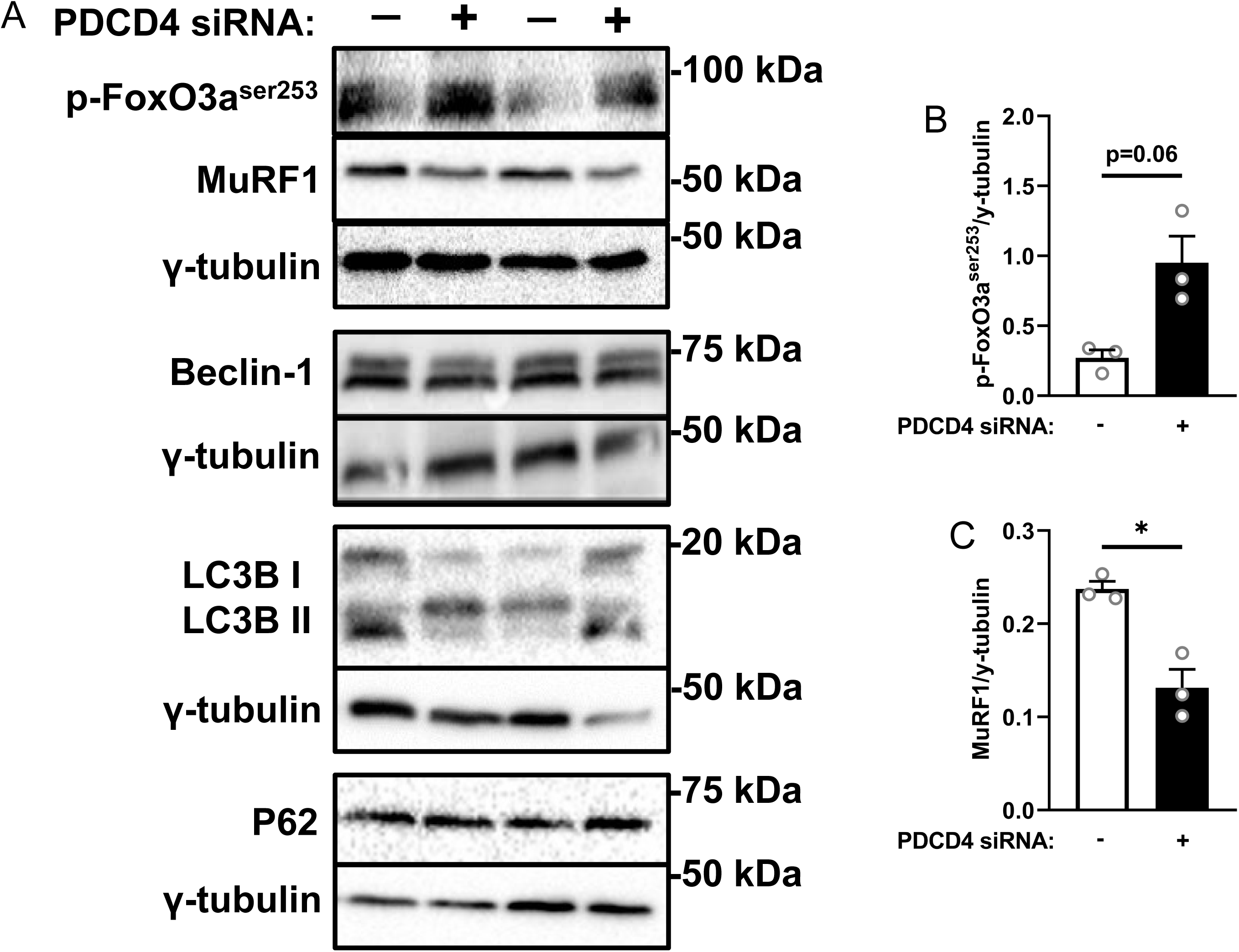
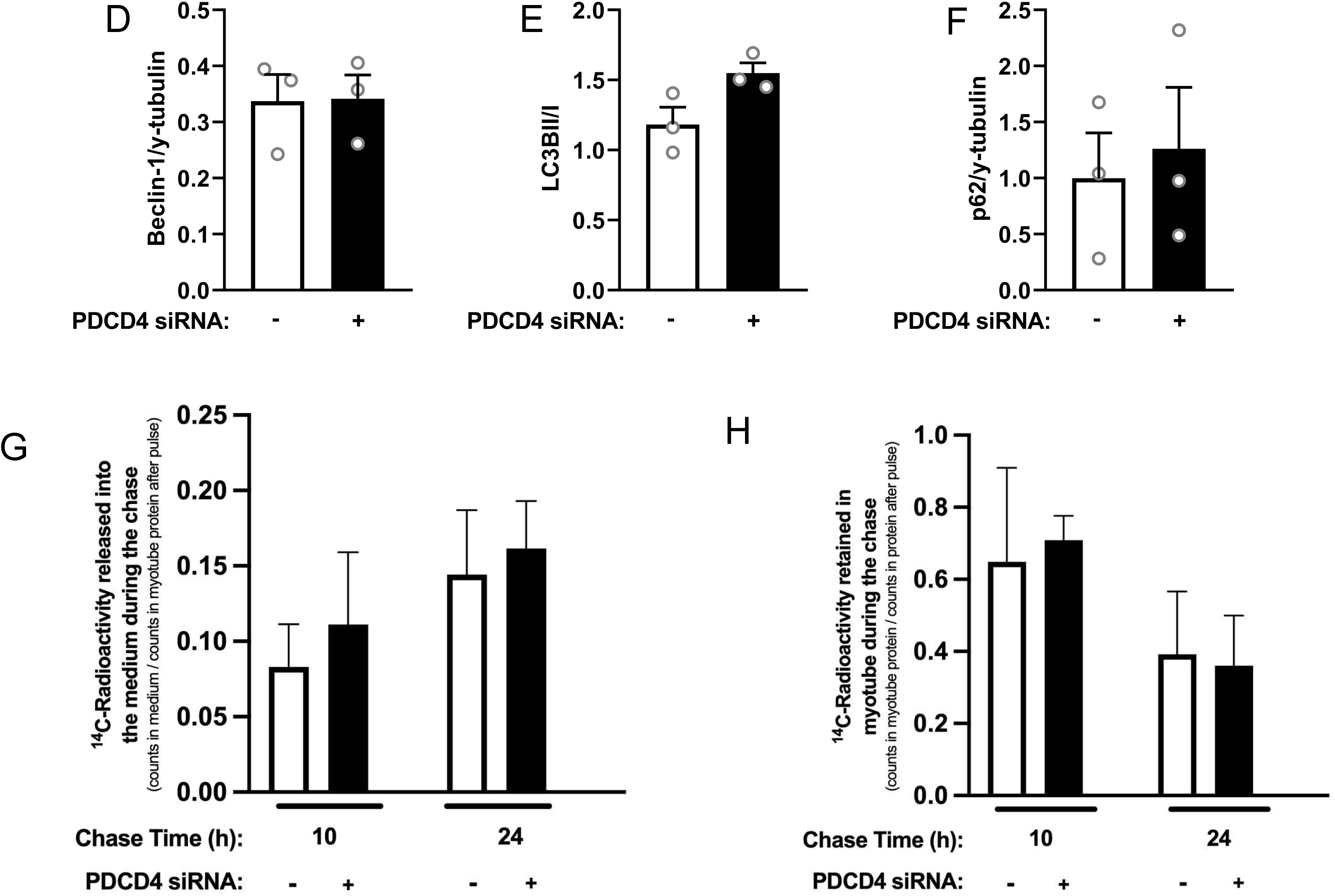
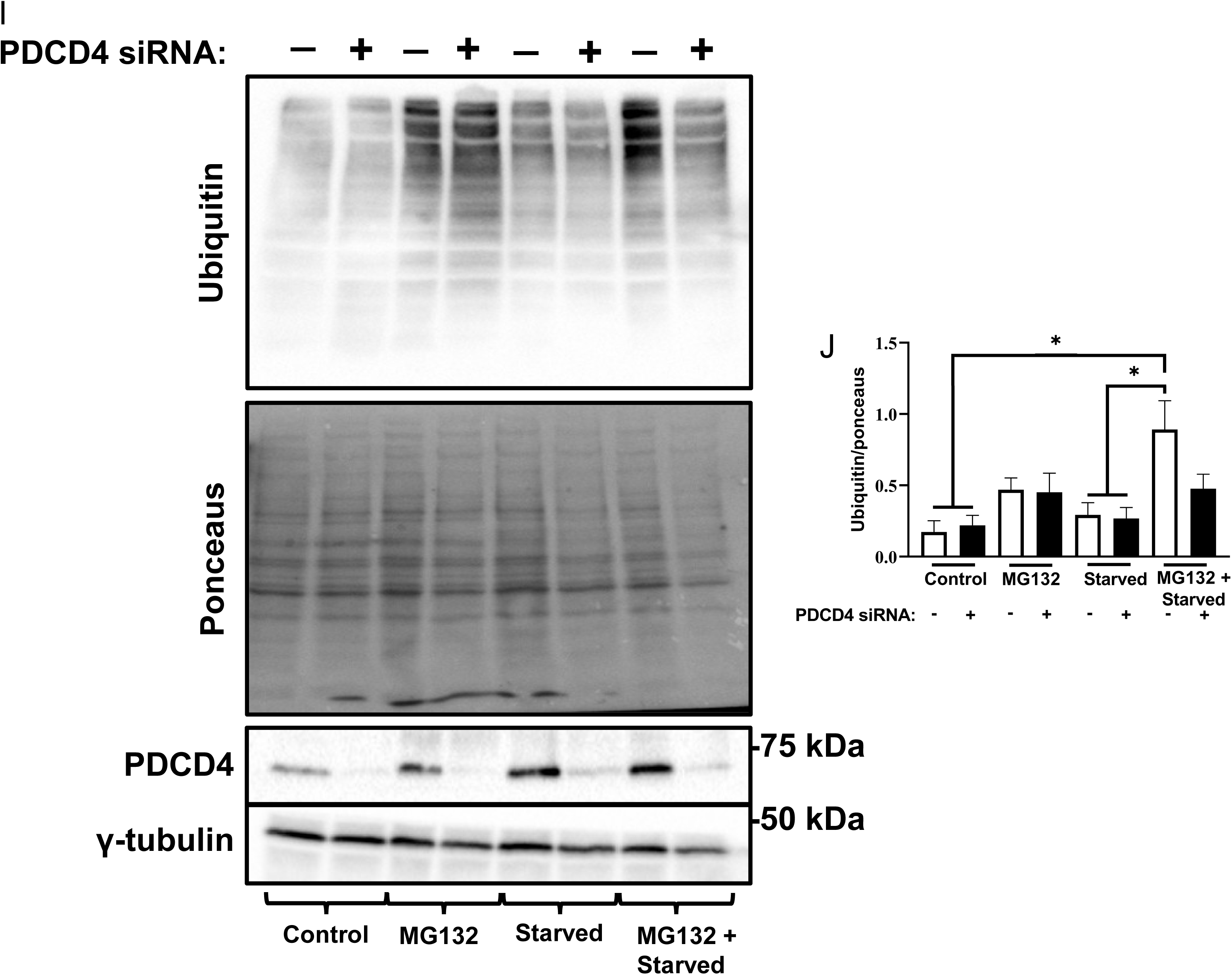
PDCD4 depletion decreases MuRF1 protein expression, but not the expression of markers of autophagy in myotubes. L6 myotubes were treated as described in figure 1. Immunoblotting **(A)** and quantified data for p-FoxO3a^ser253^ **(B)**, MuRF1 **(C)**, Beclin-1 **(D)**, LC3BII/I **(E)** and p62 **(F)** are shown. **G-H:** L6 myotubes were transfected on day 4 of differentiation and pulsed with [¹□C]-valine for 48 h to label intracellular protein pools. After the pulse, cells were chased in non-radioactive medium for either 10 h or 24 h. [¹□C]-valine radioactivity released into the incubation media during the chase period (G) and retained in cellular proteins (H) were measured by scintillation counting. For both (G) and (H), values were normalized to radioactivity measured in cellular proteins at the end of the 48-h pulse period (0□h time point), which represents the total labeled protein pool prior to the chase. **I-J.** Scrambled and PDCD4-depleted myotubes were cultured in regular differentiation medium (control), or starved for 24h with or without incubation with the proteasome inhibitor MG132 (5μM) in the last 3h of incubation. Cells were harvested and immunoblotted for ubiquitin. All data are mean ± SEM, n=3 independent experiments, with 3 technical replicates per experiment * p < 0.05.

### PDCD4-depletion does not increase myotube contraction but reduces intracellular ATP level

We wondered if the increase in the accumulation of myofibrillar proteins and the increase in myotube diameter in PDCD4-depleted myotubes conferred functional benefits. We measured sarcoplasmic reticulum (SR) Ca^2+^ release during C2C12 myotube contraction using electrical stimulation and Indo-1 AM fluorescence imaging. There was no significant difference in raw or normalized tetanic [Ca² □] □between PDCD4-depleted and control myotubes (Fig. 5A-C).

**Fig 5.**
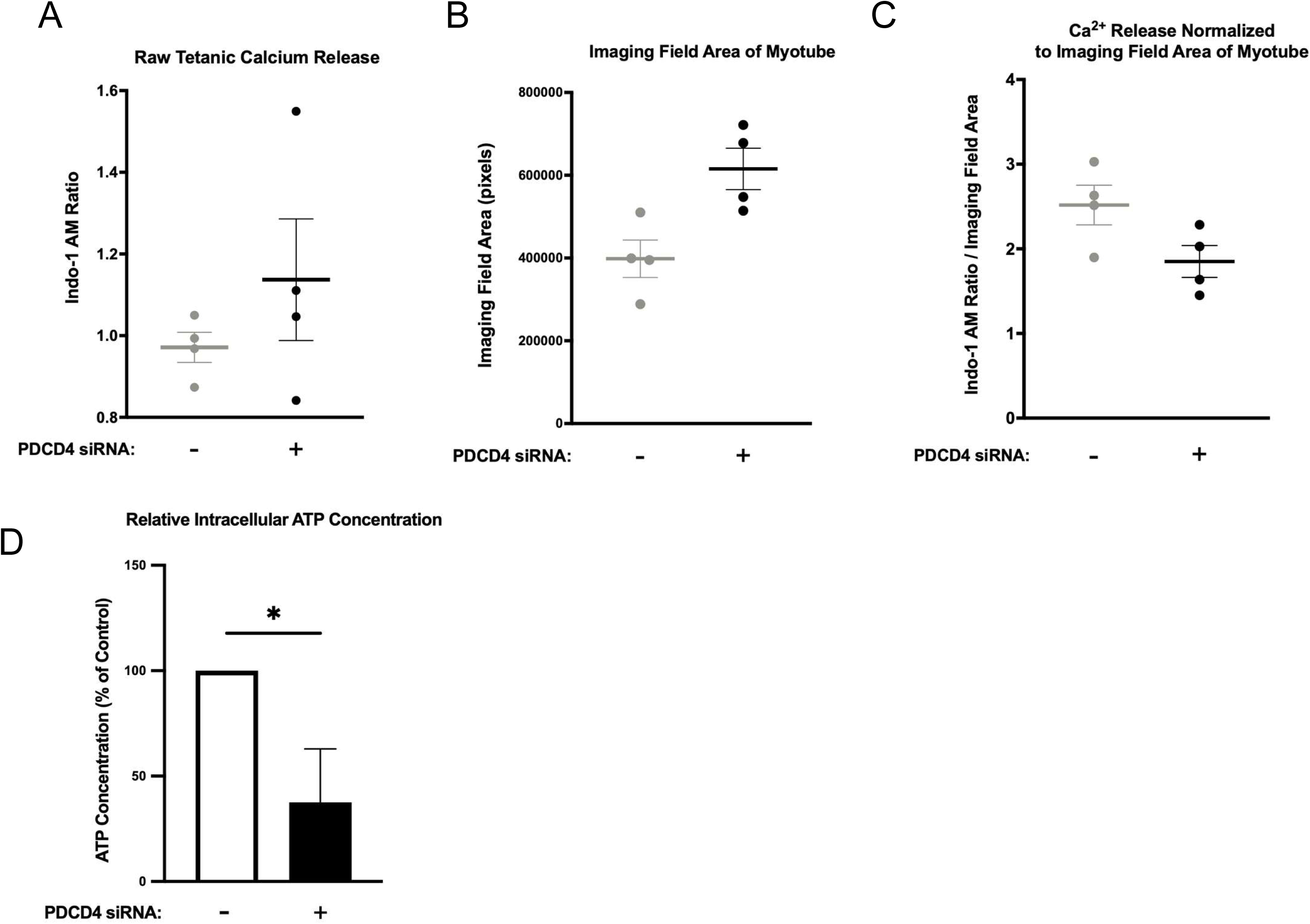
PDCD4 knockdown does not alter tetanic Ca^2+^ release in C2C12 myotubes. C2C12 myotubes were transfected with either scrambled (SCR, grey dots, ‘−’) or PDCD4 (black dots, ‘+’) siRNA oligonucleotides on d 4 of differentiation. Two days later, myotubes were electrically stimulated at 200 mA for 300 ms at 100 Hz. **A** Tetanic [Ca²□]□ was measured as the mean Indo-1 AM fluorescence ratio during 100 Hz tetanic stimulation trains. **B** The size/area of the myotube(s) in the field of view was determined by exporting a screenshot of the viewed area to ImageJ, where the area was calculated. **C** [Ca²□]□ was calculated per area of the myotube(s) in the area. Data are presented as mean ± SEM, n = 4 independent experiments. **D**. Two d after transfection with control (scrambled) or PDCD4 siRNA oligonucleotides, ATP concentration was measured using the Thermo Fisher ATP Determination Kit and normalized to total protein (µmol/g protein). To facilitate comparison, ATP levels in control myotubes were set to 100%, and ATP concentrations in PDCD4-depleted myotubes were expressed as a percentage relative to the control. Data are presented as mean ± SEM, n = 4 independent experiments, with at least 3 technical replicates per experiment, *p < 0.05.

Although anabolic signaling is elevated in PDCD4-depleted myotubes, protein synthesis was suppressed. Combined with the fact that increased myotube diameter in these myotubes did not conferring functional benefit, we wondered if these observations are related to the energy status of the myotubes. A reduction in cellular ATP availability may limit the capacity for protein synthesis despite the activation of anabolic signaling. ATP concentration was lower (-62.5%) in PDCD4-depleted myotubes when compared to the myotubes transfected with scrambled siRNA oligonucleotides (Fig. 5D).

### Reduced intracellular amino acid levels in PDCD4-depleted myotubes

Reduction in protein synthesis may also be related to altered amino acid availability. As can be seen in Fig 6A and B, 48 h after transfection, compared to control myotubes, PDCD4-depleted myotubes had reduced intracellular concentrations of many amino acids, especially the branched-chain amino acids (valine, leucine, isoleucine), proline, and tyrosine.

**Fig 6.**
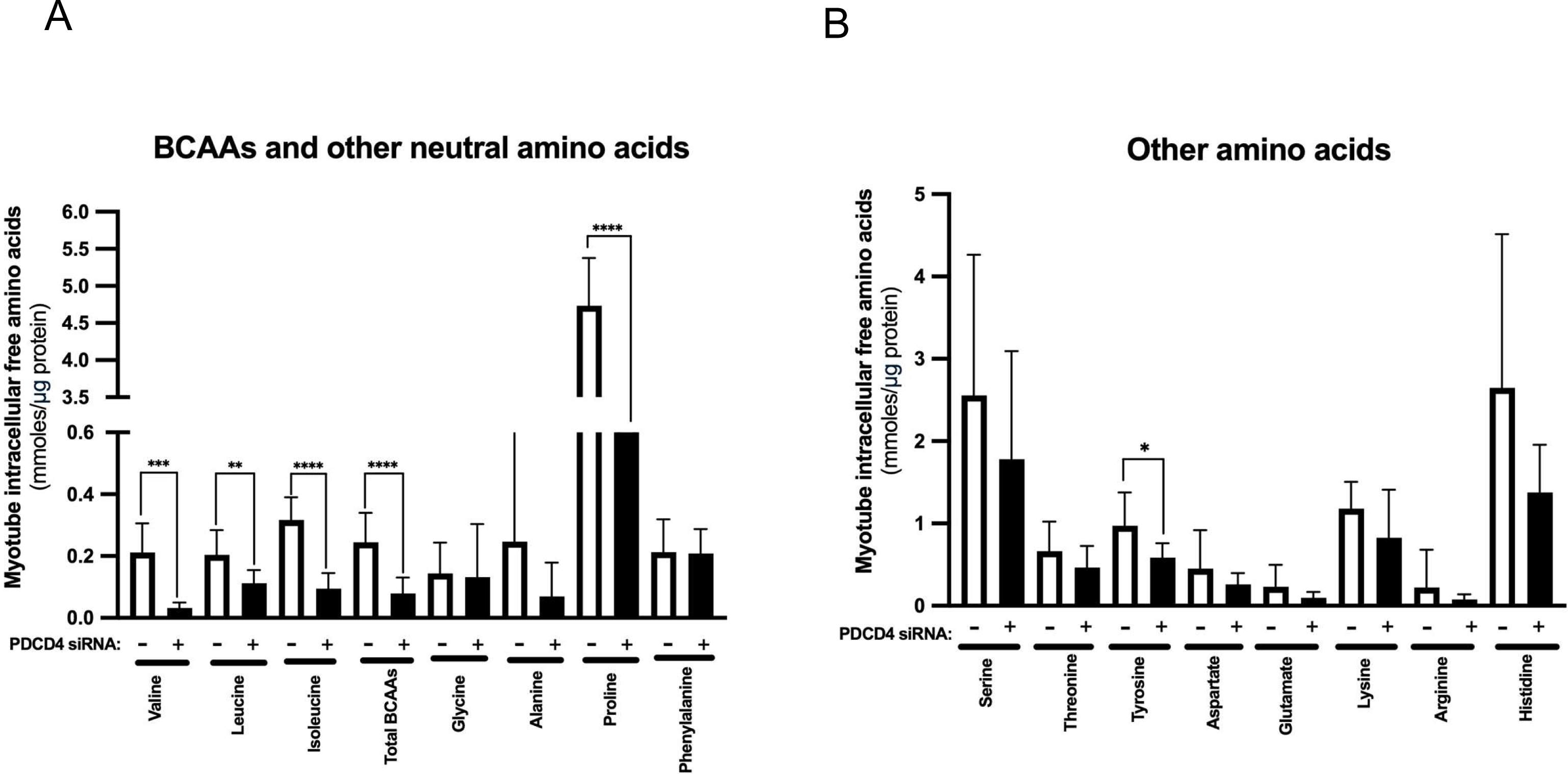
Reductions in intracellular free branched-chain amino acid levels in PDCD4-depleted myotubes. L6 myotubes were treated as described in Fig 1. Two d after transfection, samples were harvested and intracellular concentrations of amino acids were measured by high-pressure liquid chromatography (HPLC). **A**. Concentrations of the branched-chain (BCAA) and other neutral amino acids. **B** Concentrations of additional amino acids. Data are mean ± SEM, n = 3 independent experiments, with three technical replicates per experiment, *p < 0.05, **p < 0.01, ***p < 0.001, ****p < 0.0001.

### Myofibrillar protein abundance in PDCD4-depleted cells is attenuated following AKT inhibition

Because of the marked increase in AKT phosphorylation in PDCD4-depleted cells and the report that AKT is activated in conditions when protein synthesis is inhibited (37), we wondered if AKT mediated the effect of PDCD4 depletion on myotube anabolism. Therefore, we treated myotubes with PDCD4 siRNA and an AKT inhibitor either alone or in tandem. Inhibiting the activation of AKT (Fig. 7A, B) had no effect on PDCD4 abundance following siRNA treatment (Fig. 5A, C). However, treatment with the AKT inhibitor attenuated the increases in only MHC-1 myofibrillar protein content following PDCD4 depletion (Fig. 7A, D–F).

**Fig 7.**
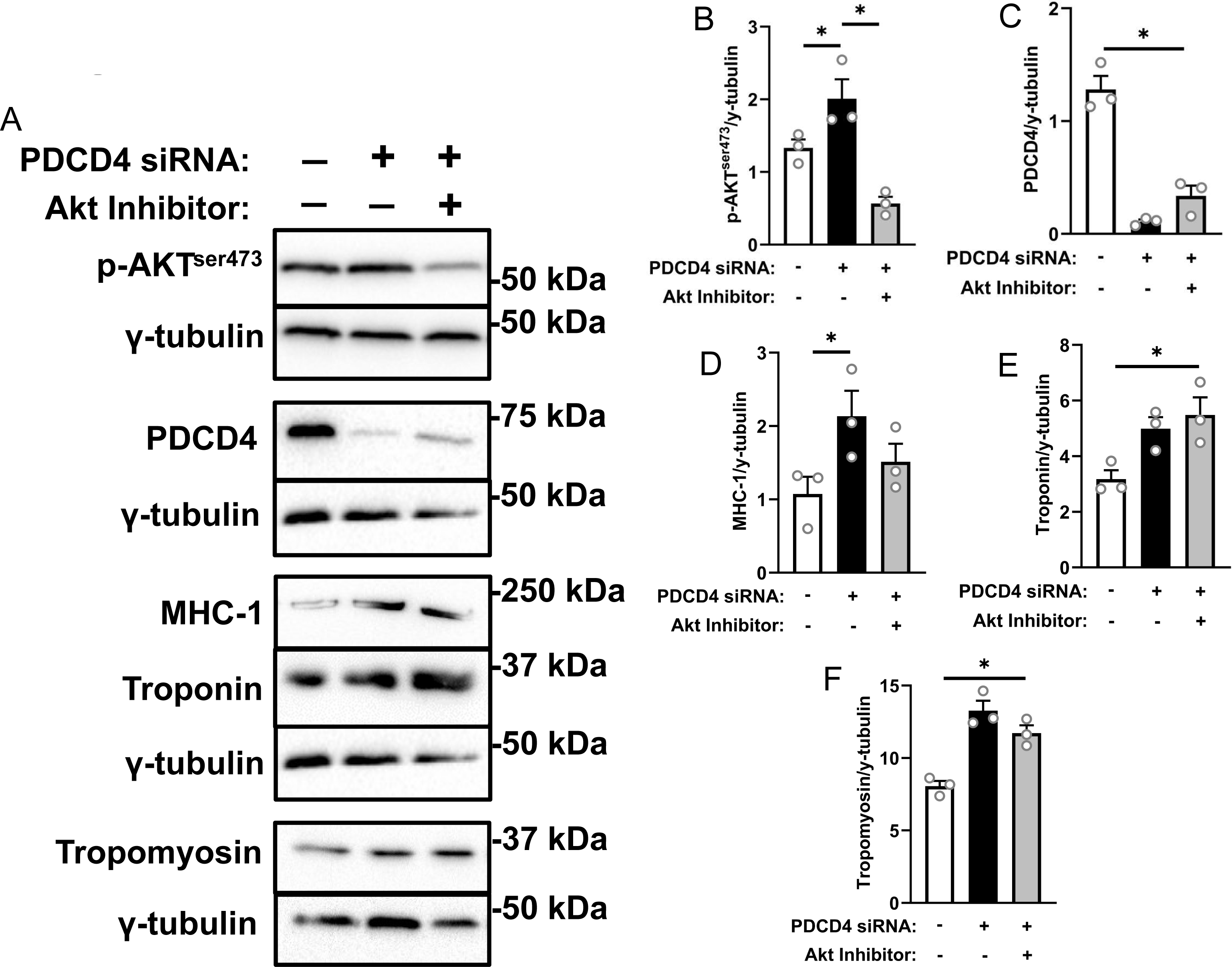
Myofibrillar protein abundance in PDCD4-depleted cells is attenuated following AKT inhibition. L6 myotubes were treated with either SCR siRNA (white bars), PDCD4 siRNA (black bars) or PDCD4 siRNA + an AKT inhibitor (grey bars). Immunoblotting **(A)** and quantified data for p-AKT^ser473^ **(B)**, PDCD4 **(C)**, MHC-1 **(D)** and troponin **(E)** and tropomyosin **(F).** Data are mean ± SEM, n=3 independent experiments, * p < 0.05.

## Discussion

The data presented in this study link PDCD4 depletion to a marked augmentation of myotube diameter and myofibrillar protein content. These changes occurred alongside an increase in the activation of some of the components of mTORC1 anabolic signaling, but not protein synthesis, likely because of reductions in intracellular amino acid levels. Although levels of MuRF1, a muscle E3 ubiquitin protein ligase, were reduced, there were no significant effects on proteolysis. Increased abundance of myofibrillar proteins did not confer any functional benefit as there was no difference in SR Ca^2+^ release in PDCD4-depleted myotubes. Mechanistically, AKT appears, at least in part, to mediate the effect of PDCD4 depletion on myofibrillar protein abundance. Combined with previous data showing that the effect of PDCD4 in cells might depend on cell type/the physiological state of the cells (26, 28), our data collectively add to the complexity in the role of PDCD4 in regulating myotube size, and metabolism.

As a tumor suppressor protein and an inhibitor of mRNA translation initiation (20), the canonical hypothesis would be for PDCD4 depletion to lead to increased protein synthesis. Indeed, this was the case for myoblasts (26, 28), but not in myotubes. In fact, protein synthesis was reduced in myotubes depleted of PDCD4. In an earlier report, PDCD4 depletion in myotubes reduced phenylalanine incorporation into total and myofibrillar proteins along with a reduction in the interactions between eukaryotic translation initiation factor 4G (eIF4G) and eIF4E (28). mTORC1/S6K1 and its substrates are more commonly studied for their effect on mRNA translation and protein synthesis, but a decrease in protein synthesis in PDCD4-depleted cells, coupled with increased myofibrillar protein abundance, suggested alterations to proteolysis. As a master regulator of anabolism, activated mTORC1 also suppresses autophagy. For example, activation of mTORC1 leads to the phosphorylation of the ATG13/ULK-1 pathway, leading to the inhibition of autophagic/lysosomal machinery (17). Thus, increased AKT and S6K1 phosphorylation observed in PDCD4-depleted cells might contribute to anabolism by suppressing autophagy. However, this was not the case in this study. Indeed, while PDCD4 depletion appeared to reduce some measures of ubiquitin-dependent proteolysis, overall proteolysis as measured by ^14^C valine pulse-chase did not reveal a significant effect of PDCD4 depletion. Because we did not examine different protein fractions (myofibrillar vs sarcoplasmic), it is possible that PDCD4-depletion had a differential impact on the synthesis/degradation of myofibrillar proteins that we did not capture, for example, by selectively increasing the synthesis of myofibrillar proteins while also suppressing the degradation of this fraction. However, in a previous study, we did not find any differences in rates of synthesis of myofibrillar vs sarcoplasmic proteins in PDCD4-depleted myotubes (28).

Alternatively, because PDCD4 is a regulator of apoptosis, it is possible that dysfunctional apoptosis that could result from PDCD4 depletion led to the accumulation of dysfunctional myotubes. Indeed, there is evidence that PDCD4 depletion leads to the development of senescent phenotype (38). This is consistent with the fact that increased myofibrillar protein abundance in PDCD4-depleted myotubes did not lead to improved SR Ca^2+^ release, where muscle contractile force is primarily regulated by cytoplasmic free [Ca^2+^] (39).

A paradoxical finding of this study is the increased AKT phosphorylation (activation) and corresponding increase in the phosphorylation of some mTORC1 targets, but reduced protein synthesis. Our data showing reduced myotube intracellular amino acid (especially the BCAA) levels and reduced ATP could explain this paradox, as amino acids and energy are needed to drive protein synthesis (40). The increase in AKT activation may also explain mTORC1 activation (as depicted by increased S6K1 phosphorylation) via the AKT-TSC1/2 signaling axis (41). As mentioned above (38), PDCD4 depletion may lead to myotube senescence. Therefore, increased mTORC1 activity in myotubes in the face of reduced amino acid level may also result from senescence, as this condition is associated with activation of mTORC1 in a manner that is insensitive to amino acid starvation (42).

Increased AKT phosphorylation leading to the phosphorylation of FoxO3a^ser253^ could explain the suppression of the ubiquitinated proteins as AKT activation would have a negative effect on MuRF1 and MAFbx/atrogin-1 expression (43). For technical reasons, we could not detect MAFbx/atrogin-1 levels, but MURF1 expression and accumulation of ubiquitinated proteins were suppressed. In a study with HEK293 and neuronal cells, inhibition of protein synthesis with two different inhibitors (cycloheximide and NSC119889) increased AKT phosphorylation (on T308 and S473), with the increase in S473 phosphorylation being much more robust. There were also increases in S6K1, S6, and FoxO3a phosphorylation (37). Apart from its effects on MuRF1 and atrogin-1 expression, AKT is involved in regulating other components of ubiquitin-dependent proteolysis. For example, it can regulate the abundance of murine double minute 2 (MDM2), the ubiquitin ligase that targets p53 for degradation (44, 45), as well as the abundance of the S-phase kinase-associated protein 2 (Skp2) (46), a protein ligase that targets many proteins for degradation (47). AKT also phosphorylates and stabilizes the deubiquitinating enzyme (DUB) ubiquitin specific proteases 8 (USP8) at Thr907 (48). Among their many roles, DUB can regulate ubiquitin dependent proteolysis by removing ubiquitin from substrates (thereby reversing the work of ubiquitin ligases) and preventing proteins from being degraded (49). These previous findings point to multi-layer regulation of proteolysis by AKT. Therefore, while some questions remain, our data along with the findings of others (37) implicate AKT in myotube anabolism in an environment of apparent protein synthesis suppression.

An intriguing question is how a reduction in the abundance of PDCD4 levels led to an increase in the phosphorylation of AKT^ser473^ and S6K1^thr389^. As the phosphorylation of AKT is responsive to diverse cellular stressors (50), the inhibition of protein synthesis seen in PDCD4-depleted cells might be the trigger that activates AKT.

In addition to PDCD4’s role in mRNA translation initiation, PDCD4 can interact/regulate the activities of other proteins, including the transcription factor AP-1, beta catenin, mitogen-activated protein kinase 1 (MAP4K1), and the basic helix-loop-helix (bHLH) transcription factor Twist (51). These mentioned proteins have roles in cell processes such as proliferation, transformation, differentiation, apoptosis, and inflammation. We did not examine these in the current study. Also, microRNA-21 (miRNA-21) negatively regulates PDCD4 level (52). On the other hand, cytoplasmic PDCD4 can also inhibit AP-1 activity, leading to reduced miRNA-21 levels. In other words, a lack of PDCD4 could increase miRNA-21 levels (53). Because miRNA-21 and other miRNAs that interact with and regulate PDCD4 expression might control the expression of targets other than PDCD4 and therefore regulate diverse cellular processes, it is possible that increased miRNAs which might result from PDCD4 depletion could affect myotube physiology and metabolism (for example, inflammation (54, 55)) at levels that we have not examined.

While we have provided possible links between reduced ATP and intracellular amino acid levels on one hand and measures of myotube contraction and protein synthesis on the other hand, it would be necessary to do a time-course study to examine which came first, and whether PDCD4 depletion impairs mitochondrial integrity (which can explain reduced ATP level). Another limitation of this study is that it was performed *in-vitro*. However, the data presented here can be leveraged to design relevant *in-vivo* studies to manipulate PDCD4 level. Because PDCD4 is a tumor suppressor protein, such *in vivo* studies should be designed in a manner that targets PDCD4 depletion in a tissue specific (skeletal muscle) manner. This will avoid undesirable consequences in tissues such as the gastrointestinal tract that are prone to PDCD4-linked cancers (56).

In conclusion, we demonstrated that depletion of PDCD4 in skeletal muscle cells enhances myotube diameter, myofibrillar protein content, and anabolic signaling but without affecting intracellular Ca^2+^ handling. These changes occurred in an apparent environment of suppression of protein synthesis, reduced intracellular ATP and amino acids levels, and minimal effects on proteolysis. Protein synthesis (57) and ubiquitin-dependent proteolysis (58) are high-energy-requiring cellular processes. Thus, the observed effects collectively suggest a coordinated response to reduced energy levels. The increase in the abundance of myofibrillar proteins are linked to AKT activation. These data point to the significance of AKT/PDCD4 axis in myotube and, by extension, potentially skeletal muscle anabolism.

## Acknowledgements

This study was funded by grants from the Natural Science and Engineering Research Council of Canada (NSERC; RGPIN-2021-03603) and from the Faculty of Health, York University, Toronto Canada to OAJA, and by grants awarded to AJC from the Natural Science and Engineering Research Council of Canada to A.J.C (RGPIN-2020-06443, DGECR-2020-00136), and the Canadian Foundation for Innovation – John R. Evan’s Leaders Fund and Ontario Research Fund awarded to A.J.C (38351).We thank Miriam Zerrouk for her assistance in some of the experiments.

## Author Contributions

S.M., and O.A.J.A. designed the study. S.M., L.A., L.D., T.A., P.N.M., and L.F. collected data and performed data analysis. S.M., L.A., L.D, T.A., L.F., A.J.C. and O.A.J.A prepared and/or edited the manuscript. All authors read and approved the final version of the manuscript.

